# Teicoplanin attenuates RNA virus infection *in vitro*

**DOI:** 10.1101/2024.09.29.615295

**Authors:** Erica Españo, Jiyeon Kim, Seong Ok Park, Bill Thaddeus Padasas, Sang-Hyun Kim, Ju-Ho Son, Jihee Oh, E-Eum Woo, Young-Ran Lee, Soon Young Hwang, Seong Kug Eo, Jeong-Ki Kim

## Abstract

Teicoplanin (TP) is a glycopeptide antibiotic used for Gram-positive bacterial infections, and it has been reported to inhibit SARS-CoV-2 and Ebola virus entry through cathepsin inhibition. Given that TP can inhibit viruses belonging to different virus families, we aimed to expand the potential targets of TP to determine whether TP can be developed as a broad-spectrum antiviral agent. Considering the original indication of TP, we first determined the effects of TP against viruses that cause respiratory tract infections and found that TP inhibits enveloped and non-enveloped RNA viruses, namely: human and avian influenza viruses; representative coronaviruses including porcine epidemic diarrhea virus (PEDV), human coronavirus OC43 (HCoV-OC43), and SARS-CoV-2; measles virus; human respiratory syncytial virus A2; and enterovirus 71 (EV-71). Representative flaviviruses, Zika virus (ZIKV) and dengue virus serotype 2 (DENV2), were also susceptible to inhibition by TP. In contrast, TP did not attenuate infection of human adenovirus 5, a non-enveloped DNA virus. Addition of TP at the endocytosis stage but not at the attachment/binding stage of PEDV infection reduced PEDV production *in vitro*, indicating cathepsin inhibition. Meanwhile, addition of TP during either the attachment/binding or the endocytosis stage of ZIKV infection reduced ZIKV particle production in host cells, and *in silico* modeling suggested that TP has potential binding pockets in the envelope proteins of ZIKV and DENV2. These results show that TP can be developed as a broad-spectrum antiviral especially against RNA viruses, with potentially different targets in the replication cycle of various viruses.

## 1 Introduction

The recent pandemic caused by severe acute respiratory syndrome coronavirus 2 (SARS-CoV-2) has highlighted the need to identify and stockpile antivirals for rapid response to future viral epidemics and pandemics. The current one-drug-one-bug approach to antiviral discovery and development, wherein drugs are targeted to a specific virus, limits drug targets to known viral threats and delays the response to future outbreaks caused by newly emerging viruses. Developing broad-spectrum antiviral agents (BSAAs) for viruses within the same family or for viruses that utilize similar infection mechanisms will be helpful in preparing for new global health threats. Direct-acting antivirals that target viral polymerases and proteases, which are typically conserved across taxonomically related viruses, are a good starting point for BSAA development. Although remdesivir and molnupiravir were originally developed for other RNA viruses, they have been approved for use against SARS-CoV-2 infection, suggesting that direct-acting antivirals can be designed against multiple targets (Eastman et al., 2020; Imran et al., 2021). Ribavirin, which is currently used for controlling hepatitis C virus (HCV) infection, has likewise shown effectivity against members of different virus families (Nyström et al., 2019; Unal et al., 2021). Other approved drugs for non-viral diseases have also been tested for potential broad-spectrum antiviral effects (e.g., hydroxychloroquine, statins, and chlorpromazine), although one is yet to be recommended for clinical application against viral infections (Mercorelli et al., 2018). This last group has the additional advantage of already established pharmacokinetic, pharmacodynamic, safety, and toxicological profiles that may accelerate emergency use authorization or licensure for future viral epidemics.

Viruses exploit host proteases at different stages of their replication cycles. Most enveloped RNA viruses that carry class I fusion proteins, including coronaviruses, influenza viruses, paramyxoviruses, pneumoviruses, and filoviruses, require proteolytic cleavage of the precursor form of the fusion protein to facilitate fusion of the virus and host membranes prior to the release of the viral genome into the host cytoplasm (Rey and Lok, 2018). Non-enveloped viruses such as human papilloma virus and reoviruses have also been shown to require cleavage of capsid proteins for viral entry and disassembly (Cerqueira et al., 2015; Ebert et al., 2002). Viral proteins may also require proteolytic processing for the maturation of non-infectious to infectious viral particles inside the cell (e.g., flaviviruses and alphaviruses) (Pierson and Diamond, 2012; Skidmore and Bradfute, 2023). Thus, targeting either viral or host proteases is now considered an effective strategy for treating viral infections, as exemplified by clinically approved antiviral regimens that include inhibitors of HIV and HCV proteases (Majerová and Konvalinka, 2022).

The cathepsin family consists of several proteases, most of which are cysteine proteases, that exhibit optimal activity in the acidic environment of lysosomes (Yadati et al., 2020). As such, they are primarily found in endo/lysosomal compartments and are involved in autophagy. Cathepsins are also found in the extracellular space and are typically needed for remodeling the extracellular matrix and repairing the plasma membrane. Among the first studies to report cathepsin involvement in viral infection was one that showed that cathepsin B (CatB) and cathepsin L (CatL) cleave Ebola virus (EBOV) glycoprotein 1 (GP1) in a multistep process (Chandran et al., 2005). CatB and CatL were also found to mediate the disassembly of reovirus particles in the endosome (Ebert et al., 2002). Different virus families appear to utilize cathepsins as part of the infection cycle in addition to other types of proteases (e.g., serine proteases), making cathepsins a viable target for antiviral development (Scarcella et al., 2022). Teicoplanin (TP), a glycopeptide antibiotic that is structurally related to vancomycin, has been observed to inhibit entry of viral particles pseudotyped with the envelope glycoproteins of EBOV, Middle Eastern respiratory syndrome coronavirus (MERS-CoV), and SARS-CoV into host cell lines (Wang et al., 2016; Zhou et al., 2016). In the same study, the effects of TP on EBOV entry were attributed to inhibition of cathepsin L (CatL). TP was also reported to inhibit human immunodeficiency virus 1 (HIV-1) *in vitro* (Balzarini et al., 2003). These studies suggest that cathepsin inhibition through TP treatment can potentially attenuate infection of several viruses that share infection mechanisms.

Given the ability of TP to attenuate infection of a number of viruses *in vitro*, we here aimed to expand the breadth of antiviral targets of TP as a potential BSAA by testing the effects of TP on human and veterinary viral pathogens belonging to different families. Taking into consideration the original indication of TP as an antibiotic, we first focused on enveloped RNA viruses that cause human respiratory tract infections that are typically associated with bacterial infections (influenza viruses, coronaviruses, paramyxovirus, and pneumovirus). In addition, we tested a non-enveloped RNA virus representative (enterovirus 71) and a non-enveloped DNA virus (human adenovirus 5), both of which can be transmitted through the respiratory route. We also tested the effects of TP on flavivirus infection *in vitro*. The results of this study support existing evidence that TP exhibits broad antiviral activity and should be considered for development into a BSAA.

## 2 Materials and Methods

### 2.1 Cells and Viruses

Lung epithelial A549 cells, mink lung Mv1Lu cells, HeLa-derived HEp-2 cells, vervet monkey kidney Vero and Vero E6 cells were grown in Minimum Essential Medium (MEM; Gibco, ThermoFisher Scientific, Waltham, MA) supplemented with 10% heat-inactivated fetal bovine serum (FBS; Gibco, Carlsbad, CA), 1% antibiotic-antimycotic solution (Gibco), and 3% _-glutamine (Gibco). Madin-Darby canine kidney (MDCK) cells were grown in MEM supplemented with 10% fetal bovine serum, 1% antibiotic-antimycotic solution, 3% _-glutamine, and 1% MEM vitamin solution (Gibco). All cell cultures (with and without virus) were incubated in a humidified atmosphere with 5% CO_2_.

Eight influenza viruses were used in this study: A/Puerto Rico/8/1934 (A/PR/8/34; H1N1); A/California/04/2009 (A/CA/04/09; H1N1); A/Philippines/2/1982 (A/PH/2/82; H3N2); A/Brisbane/10/2007 (A/Bris/10/07; H3N2); A/Aquatic Bird/Korea/CN2/2009 (A/AB/Kor/CN2/09; H5N2); A/Aquatic Bird/Korea/CN5/2009 (A/AB/Kor/CN5/09; H6N5); A/Chicken/Korea/01310/2001 (A/Ck/Kor/01310/01; H9N2); and B/Seoul/32/2011 (B/Seoul/32/11; Yamagata-like). All influenza viruses were propagated in MDCK cells in MEM + BSA, consisting of MEM supplemented with 0.3% bovine serum albumin (Sigma-Aldrich, St. Louis, MO), 1% antibiotic-antimycotic solution, 3% _- glutamine, 1% MEM vitamin solution and freshly added 1.0 μg/ml tosylsulfonyl phenylalanyl chloromethyl ketone (TPCK)-trypsin (Worthington Biochemical Corporation, Lakewood, NJ) incubated at 37 °C. After incubation for 72 h, the supernatants of infected MDCK cells were harvested, clarified through centrifugation, and stored in aliquots at –80 °C until use. All other viruses were harvested and stored similarly. The assay medium for influenza viruses was the same medium used for influenza virus propagation.

Zika virus (ZIKV) PRVABC59 strain (ATCC VR-1843) was propagated in Vero cells with MEM + BSA without TPCK-trypsin. ZIKV was harvested after 5 days of incubation at 37 °C. Dengue virus serotype 2 (DENV2; ATCC VR-1584) was propagated in Vero E6 cells in MEM + 1% FBS consisting of 1× MEM, 1% heat-inactivated FBS, 1% antibiotic-antimycotic solution, 3% _-glutamine, and 1% MEM vitamin solution for 6 days at 37 °C. Measles virus (Edmonston strain; ATCC VR-24) was propagated in Vero cells in MEM + 1% FBS for 7 days at 37 °C. Human RSV-A2 (HRSV-A2) was kindly provided by the International Vaccine Institute (Seoul, South Korea) and was propagated in HEp-2 cells in MEM + 1% FBS for 5 days at 37 °C. Human coronavirus OC43 (HCoV-OC43; ATCC VR-1558) was propagated in Mv1Lu cells in MEM + 1% FBS for 7 days at 37 °C. Human adenovirus 5 (HAd5; ATCC VR-5) was propagated in A549 cells in MEM + 1% FBS for 3 days at 37 °C. Enterovirus 71 (EV-71; ATCC VR-1432) was propagated in Vero cells in MEM + BSA without TPCK-trypsin for 4 days at 37 °C. For all these viruses, the assay medium was the same as the propagation medium.

Cell-culture adapted porcine epidemic diarrhea virus (PEDV; DR13 strain) was propagated in Vero cells in medium consisting of Dulbecco’s modified Eagle medium (DMEM) with sodium pyruvate supplemented with 0.02% yeast extract, 0.3% tryptose phosphate broth, 1% antibiotic-antimycotic solution, and freshly added trypsin (5 µg/mL) (Song, 2003). The infected cells were incubated for 66 hours at 37 °C. The assay medium for PEDV in Vero cells was MEM + BSA without TPCK-trypsin.

SARS-CoV-2 (Wuhan prototype; NCCP43330) was provided by the National Culture Collection for Pathogens in South Korea and propagated in Vero E6 (ATCC CRL-1586) with 2% FBS-supplemented Dulbecco’s Modified Eagle Medium (DMEM). Virus titration was performed using Vero E6 with the cytopathic effect and standard plaque assays. All experiments with SARS-CoV-2 were performed in biosafety level 3 (BSL3) facilities at Core Facility Center for Zoonosis Research, Jeonbuk National University (Iksan, South Korea).

### 2.2 TP preparation and treatment of cells

TP powder was kindly provided by the Institute of Pharmaceutical Technology, Dongkook Pharmaceutical Co. Ltd. (Jincheon-gun, Chungbuk, South Korea) and was stored at 4 °C. Pre-weighed aliquots of TP powder to masses appropriate for subsequent assays were prepared and stored at 4 °C until the day of use. Several aliquots of TP powder were prepared on the same batch to reduce inter-assay variability due to weighing.

On the day of the antiviral assay, a pre-weighed TP aliquot was dissolved in the assay medium appropriate to the virus for testing, and, immediately before the assay, TP was serially diluted in the assay medium to 2× of the required concentrations. Then, the prepared 2× dilutions of TP in assay medium (50 µL/well) were added to infected cells (n = 3 per dilution). Untreated, infected controls were prepared by adding assay medium to infected cells (n = 3). Untreated, uninfected controls were prepared by adding assay medium to non-infected cells on the same plate (n = 3). Cytotoxicity assays of TP on various cell lines were similarly performed using high concentrations of TP added to uninfected cells (n = 3 per dilution). For determining cell viability, at the end of the incubation time required for each virus, the old assay medium was discarded and replaced with fresh assay medium (100 µL/well), to which 10 µL/well of EZ-Cytox (DoGenBio; Seoul, South Korea) was added. The cells were incubated for 3–4 hours at 37 °C in a humidified atmosphere with 5% CO_2_, and absorbance at 450 nm (Abs_450_) was determined using a microplate reader. Assays were repeated at least twice for calculations of the relative half-maximal effective (EC_50_) and cytotoxic (CC_50_) concentrations. All TP preparations in the assay medium were used on the same day.

### 2.3 Determining the effects of TP on influenza virus-induced cytopathic effects (CPE)

MDCK cells were seeded into 96-well plates at a density of 2.0 × 10^4^ cells/well in the appropriate growing medium and incubated overnight at 37 °C. The following day, the cells were washed twice with PBS and then infected with the following influenza viruses (50 µL/well): 1,000 TCID_50_^/^well of A/PR/8/34 (H1N1), A/CA/04/09 (H1N1), A/PH/2/82 (H3N2), A/Bris/10/07 (H3N2), A/AB/Kor/CN5/09 (H6N5), A/AB/Kor/CN2/09 (H5N2), A/Ck/Kor/01310/01 (H9N2) or 100 TCID_50_^/^well of B/Seoul/32/11. The cells were treated with TP as indicated in Section 2.2, and the MDCK cells were incubated for 3 days at 37 °C prior to determination of cell viability as in Section 2.2.

### 2.4 Determining the effects of TP on CPE induced by coronaviruses

For PEDV infection, Vero cells were seeded into 96-well plates at a density of 1.5 × 10^4^ cells/well in the appropriate growing medium and incubated overnight at 37 °C. The cells were washed once with PBS and then infected with PEDV in assay medium (50 µL/well) at a multiplicity of infection (MOI) of 0.001. The next day, cells were treated with TP and were incubated for ∼66 hours at 37 °C. For HCoV-OC43 infection, Mv1Lu cells were seeded into 96-well plates at a density of 2.5× 10^4^ cells/well in the appropriate growing medium and incubated overnight at 37 °C in a humidified chamber with 5% CO_2._ The following day, the cells were washed twice with PBS and then infected with HCoV-OC43 (200 TCID_50_/well; 50 µL/well) in assay medium. The cells were treated with TP and then incubated for 7 days at 37 °C.

For SARS-CoV-2 infection, Vero E6 cells seeded in 96-well plates were washed once with FBS-free DMEM and then infected with SARS-CoV-2 (100 TCID_50_/well; 50 µL) in FBS-free DMEM. Following 1 h incubation for virus absorption, the virus inoculum was removed, and Vero E6 cells were treated with 2% FBS-supplemented DMEM containing TP at the desired concentrations. The Vero E6 cells were then incubated until CPE was seen in the virus-infected controls. The cells were stained with for 5 min with 0.2% crystal violet solution in MeOH. After removing crystal violet, the plates were washed with tap water and fully dried at room temperature. Crystal violet was dissolved in MeOH, and the optical density (OD) of each well was measured at 570 nm. The extent of CPE in each well was calculated by comparing the OD values of the treated cells with those of the virus control wells.

### 2.5 Determining the effects of TP on CPE induced by measles virus and HRSV-A2

For measles virus infection, Vero cells were seeded into 96-well plates at a density of 1.0 × 10^4^ cells/well in the appropriate growing medium and incubated overnight at 37 °C in a humidified chamber with 5% CO_2._ The cells were washed once with PBS and then infected with measles virus in assay medium (500 TCID_50_/well; 50 µL/well). The cells were treated with TP and were incubated for 7 days at 37 °C. For infection with HRSV-A2, HEp-2 cells were seeded into 96-well plates at a density of 2.0 × 10^4^ cells/well. The cells were washed once with PBS and then infected with HRSV-A2 (70 TCID_50_/well; 50 µL/well) in assay medium. The cells were treated with TP and incubated for 7 days at 37 °C.

### 2.6 Determining the effects of TP on CPE induced by EV-71 and HAd5

For EV-71 infection, Vero cells were seeded into 96-well plates at a density of 1.2 × 10^4^ cells/well in the appropriate growing medium and incubated overnight at 37 °C. The next day, the cells were washed once with PBS and then infected with EV-71 in assay medium (500 TCID_50_/well; 50 µL/well). The cells were treated with TP and incubated for 4 days at 37 °C in a humidified atmosphere with 5% CO_2_. For HAd5 infection, A549 cells were seeded into 96-well plates at a density of 3.5 × 10^4^ cells/well in the appropriate growing medium and incubated overnight at 37 °C in a humidified chamber with 5% CO_2._ The next day, the cells were washed once with PBS and then infected with HAd5 in assay medium (4,000 TCID_50_/well; 50 µL/well).

### 2.7 Determining the effects of TP on CPE induced by Zika virus and dengue virus serotype 2

For ZIKV infection, Vero cells were seeded into 96-well plates at a density of 1.2 × 10^4^ cells/well in the appropriate growing medium and incubated overnight at 37 °C. The next day, the cells were washed once with PBS and then infected with ZIKV in assay medium (MOI of 0.02/well; 50 µL/well). The cells were treated with TP and incubated for 5 days at 37 °C. For DENV2 infection, Vero E6 cells were seeded into 96-well plates at a density of 2.5 × 10^4^ cells/well in the appropriate growing medium and incubated overnight at 37 °C. The cells were washed once and then infected with DENV2 (MOI of 0.02/well; 50 µL/well). The cells were treated with TP and incubated for 5 days at 37 °C.

### 2.8 Determining % CPE inhibition, EC_50_, and CC_50_

Percent cell viability (%CV) of the cells following treatment and infection were determined by dividing the Abs_450_ of the treated cells by the mean Abs_450_ of the untreated, uninfected cell (uninfected controls) multiplied by 100. Based on the %CV values, % CPE reduction was then calculated using the following formula:

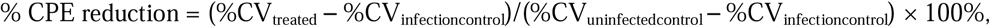

where %CV_treated_ is the %CV of the treated well, %CV_infectioncontrol_ is the mean %CV of the untreated, infected cell control group; and %CV_uninfectedcontrol_ is the mean %CV of the untreated, uninfected cell control group.

For relative EC_50_ calculations, the %CV_treated_ values were first normalized relative to the maximum and minimum %CV_treated_ in the dataset, where the minimum %CV _treated_ was equivalent to the %CV_infectioncontrol_. The means of the normalized values per assay were then used as datapoints to determine the mean relative EC_50_ through the “log(inhibitor) vs. normalized response – Variable slope” mode under the nonlinear regression analysis function of GraphPad Prism 10 (GraphPad Software, Boston, MA). For relative CC_50_ calculations, the %CV_treated_ values were first normalized using 100% (equivalent to %CV_uninfectedcontrol_) as maximum and the lowest %CV_treated_ in the dataset as the minimum. The means of the normalized values per assay were then used to determine the mean relative CC_50_ through the “log(inhibitor) vs. normalized response – Variable slope” mode under the nonlinear regression analysis function of GraphPad Prism 10. All EC_50_ and CC_50_ values were obtained from 2 to 3 independent assays.

### 2.9 Determining the effects of TP on PEDV growth kinetics

Overnight Vero cell cultures (6.0 × 10^6^ cells/well) were prepared in 6-well plates. The cells were washed once with PBS. TP (200 μM) or CQ (10 μM; positive control) in PEDV assay medium was added to the cell monolayer along with PEDV (MOI of 0.01) in assay medium for a total volume of 2 mL per well (n = 3 per treatment). Untreated controls (PEDV only, n = 3) and uninfected, treated controls (n = 1) were similarly prepared. Supernatant from the infected groups were collected at 1, 6, 12, 18, 24, 30, and 36 hours post-infection (hpi), clarified through centrifugation, and stored at –80 °C until subsequent plaque assay.

For determining viral titers through plaque assay, Vero cells (2.0 × 10^5^ cells per well) were prepared in 12-well plates two days before the assay. On the day of the assay, PEDV infection supernatants were serially diluted 10-fold (10^-1^–10^-5^) in PEDV assay medium. The Vero cells were washed once with PBS and infected with 250 μL of the serially diluted supernatants for 1.5 h at 37 °C. The inocula were removed, and the plaque overlay (0.9% bacteriological agar in PEDV assay medium) was added to each well. After 3 days, the plaque overlays were removed, and the plaques were stained and fixed with a solution of 0.1% crystal violet and 10% formaldehyde. The mean PFU/mL for each treatment was calculated (n = 3 per treatment).

### 2.10 Determining the effects of TP on PEDV and ZIKV early entry stages

For determining the effects of adding TP during the attachment/binding stage of PEDV and ZIKV infection, Vero cell cultures were prepared in 6-well plates (3.5 × 10^5^ cells/well) two days before the assay (until 95% confluence). On the day of the assay, the cells were washed once with PBS. For PEDV, the cells were infected with 200 PFU of PEDV per well in the presence of 200 μM TP, 10 μM CQ or assay medium (1 mL total volume) for 2 h at 4 °C. For ZIKV, the cells were infected with 400 plaque-forming units (PFU) of ZIKV per well in the presence of 400 μM TP, 800 μM TP, 25 μM CQ, or assay medium (1 mL total volume) for 2 h at 4 °C. The inoculum + drug mixture was removed, and the cells were washed twice with ice-cold PBS to remove unbound virus particles. The plaque overlay was added (3 mL/well), and the cells were incubated at 37 °C for 3 (PEDV) or 4 (ZIKV) days prior to staining. Percent plaque production from each treatment was calculated as the average number of plaques (PFU) of the treatment group divided by the average number of plaques formed in the untreated group multiplied by 100 (n = 3 per treatment).

For determining the effects of adding TP during the endocytosis stage of PEDV and ZIKV infection, Vero cell cultures were prepared in 6-well plates (3.5 × 10^5^ cells/well) two days before the assay (until 95% confluence). On the day of the assay, the cells were washed once with PBS. For PEDV, the cells were infected with PEDV (200 PFU in 1 mL assay medium per well) at 4 °C for 2 h. The inoculum was removed, and the cells were washed twice with ice-cold PBS. Then, the cells were treated with 200 μM TP, 10 μM CQ, or medium only (1 mL in assay medium) for 2 h at 37 °C. For ZIKV, the cells were infected with 400 PFU (in 1 mL assay medium per well), and then incubated at 4 °C for 2 h. The inoculum was removed, and the cells were washed twice with ice-cold PBS. Then, the cells were treated with 400 μM TP, 800 μM TP, 25 μM CQ, or assay medium only (1 mL in assay medium) for 2 h at 37 °C. For both viruses, after incubation at 37 °C, the compounds were removed, and the cells were washed once with PBS and treated with 300 μL citric acid buffer (40 mM citric acid, 10 mM KCl, 135 mM NaCl; pH 3.0) for 30 seconds, and then washed again with PBS. The plaque overlay was added (3 mL/well), and the cells were incubated at 37 °C for 3 (PEDV) or 4 (ZIKV) days prior to staining. Fixing and staining were all performed as in Section 2.9. Percent plaque production from each treatment was calculated as the average number of plaques (PFU) of the treatment group divided by the average number of plaques formed in the untreated group multiplied by 100 (n = 3 per treatment).

### 2.11 Predicting binding sites of TP on ZIKV and DENV2 envelope proteins

The 3D structure of TP was obtained from ChemSpider (ID: **17288587**), and energy minimization was performed using the default values in Open Babel included in PyRx ver. 0.8 (Dallakyan and Olson, 2015). Then, AutoDock Vina in the PyRx package was used to predict the docking sites of TP on the ZIKV (PDB: **5JHM**) and DENV2 (PDB: **1OKE**) envelope (E) proteins. The top model (lowest binding affinity) was selected. Discovery Studio Visualizer ver. 21.1 (Biovia Dassault Systèmes, Vélizy-Villacoublay, France) was used to view the results of the docking prediction and to determine amino acid residues predicted to form bonds with TP.

### 2.12 Statistical analysis

Student’s *t*-test was used to determine significant differences between pairs of groups (*p*_<_0.05).

## 3 Results

### 3.1 Teicoplanin attenuates influenza virus-induced CPE

Like vancomycin, TP is primarily used as antibiotic against infections with Gram-positive bacteria, including methicillin-resistant *Staphylococcus aureu*s (Svetitsky et al., 2009). Late stages of viral respiratory infections are correlated with bacterial infections termed as either co-infections (concurrent with viral infection) or superinfections (subsequent to viral infection), with *S. aureus* and *Streptococcus pneumoniae* accounting for the majority of such cases (Oliva and Terrier, 2021). Considering the original indication of TP, we first tested whether TP could inhibit the infection of viruses that cause respiratory tract infections. Among these were influenza viruses, which are the viruses most commonly associated with secondary bacterial infections (Oliva and Terrier, 2021).

Our CPE-based assay showed that infection of MDCK cells with all the influenza A viruses we tested were attenuated by TP in a dose-dependent manner at non-cytotoxic concentrations of TP (Fig. 1; Supplementary Fig. S1). However, viruses had varied susceptibilities to TP as indicated by the maximum inhibitory effects of TP (Fig. 1) and the EC_50_ values of TP (Table 1) against the different subtypes and strains of influenza A virus. Both H1N1 strains (A/PR/8/34 and A/CA/04/09) that we tested were highly susceptible to inhibition by TP, with TP able to rescue MDCK cells completely albeit at relatively high concentrations. Both oseltamivir-susceptible (A/PH/2/82) and oseltamivir-resistant (A/Brisbane/10/07) H3N2 viruses were also susceptible to TP treatment at concentrations slightly lower than those needed for H1N1 inhibition (Song et al., 2021). Notably, the avian influenza virus isolates we tested (A/H5N2, A/H6N5, and A/H9N2) (Nam et al., 2011; Nam et al., 2017) were all inhibited by TP, with the avian A/H9N2 isolate being the most susceptible, having the lowest EC_50_ and with 100% CPE reduction. The influenza B virus (B/Seoul/32/11; Yamagata-like) we tested was also susceptible to TP treatment; however, it seemed to be the least susceptible to TP among all the influenza viruses we tested based on the 71% maximal CPE reduction following TP treatment. Overall, both influenza A and B viruses appear to be susceptible to inhibition by TP.

**Table 1.** Half-maximal effective concentrations of teicoplanin against CPE induced by viruses belonging to different families.

| Family | Virus | Host cell | Maximum CPE reduction | <sup>a</sup>EC<sub>50</sub> (μM) | CC<sub>50</sub> (mM) | SI |
| --- | --- | --- | --- | --- | --- | --- |
| *Adenoviridae* | HAd5 | A549 | 0% | No activity | >0.5 | - |
| *Coronaviridae* | PEDV | Vero | 100% | 60.0 | 7.3 | 121.7 |
|  | HCoV-OC43 | Mv1Lu | 56% | 33.6 | 193.2 | 5.8 |
|  | SARS-CoV-2 | Vero | 80% | 55.5 | 7.3 | 131.0 |
| *Flaviviridae* | DENV2 | Vero E6 | 100% | 68.3 | >0.38 | >5.6 |
|  | ZIKV | Vero | 100% | 192.5 | 7.3 | 37.9 |
| *Orthomyxoviridae* | A/PR/8/34 (H1N1) | MDCK | 91% | 200.1 | 1.7 | 8.5 |
|  | A/CA/04/09 (H1N1) |  | 100% | 363.0 |  | 4.7 |
|  | A/PH/2/82 (H3N2) |  | 79% | 210.0 |  | 8.1 |
|  | A/Brisbane/10/07 (H3N2) |  | 91% | 195.3 |  | 8.7 |
|  | A/Kor/CN2/09 (H5N2) |  | 91% | 217.2 |  | 7.8 |
|  | A/Kor/CN5/09 (H6N5) |  | 86% | 282.7 |  | 6.0 |
|  | A/Kor/01310/01 (H9N2) |  | 100% | 101.9 |  | 16.7 |
|  | B/Seoul/32/11 (Yamagata-like) |  | 71% | 145.8 |  | 11.7 |
| *Paramyxoviridae* | Measles | Vero | 68% | 43.74 | 7.3 | 166.9 |
| *Picornaviridae* | EV71 | Vero | 60% | 84.6 | 7.3 | 86.3 |
| *Pneumoviridae* | HRSV-A2 | HEp-2 | 100% | 84.0 | >1.0 | >11.9 |

<sup>a</sup>EC<sub>50</sub> values were calculated relative to the minimum and maximum % cell viability and were calculated based on the means of at least two independent assays.
SI: selectivity index (CC<sub>50</sub>/EC<sub>50</sub>)

**Fig. 1.**
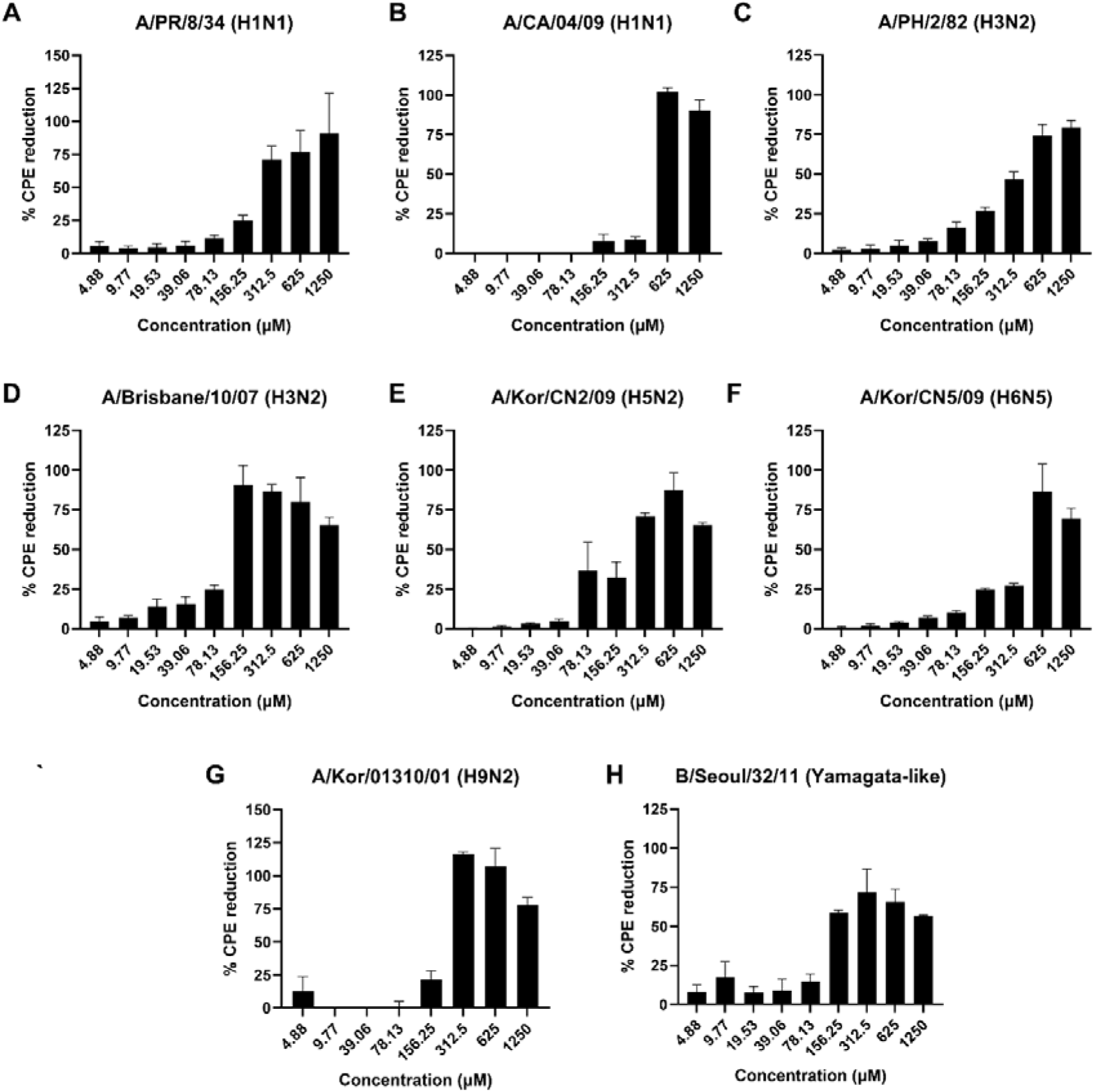
Effects of teicoplanin (TP) on cytopathic effects (CPE) induced by influenza viruses. Indicated concentrations of TP in influenza virus assay medium were prepared and used to treat Madin-Darby canine kidney (MDCK) cells infected with 1,000 TCID_50_^/^well of **A,** A/PR/8/34 (H1N1); **B,** A/CA/04/09 (H1N1); **C,** A/PH/2/82 (H3N2); **D**, A/Bris/10/07 (H3N2); **E,** A/AB/Kor/CN2/09 (H5N2); **F,** A/AB/Kor/CN5/09 (H6N5); **G,** A/Ck/Kor/01310/01 (H9N2); or **H,** 100 TCID_50_^/^well of B/Seoul/32/11 (Yamagata-like). Percent cytopathic effect reduction (% CPE reduction) was calculated relative to the viability of the untreated infected and uninfected controls (Ctrl). Data show means ± SEM; n = 3 per TP concentration. Each graph represents one of at least two independent assays.

### 3.2 Teicoplanin reduces coronavirus-induced cytopathic effects

Considering the inhibitory effects of TP on influenza viruses, we next tested whether TP could also attenuate infection with various coronaviruses. A previous study has already reported that TP was able to inhibit SARS-CoV and MERS-CoV pseudovirus entry *in vitro* (Zhou et al., 2016). Corroborating the results of this study, we saw that PEDV, a representative Alphacoronavirus, and human coronavirus OC43 (HCoV-OC43), a representative common cold Betacoronavirus, were both susceptible to inhibition by TP in a dose-dependent manner at non-cytotoxic concentrations, with TP rescuing PEDV-infected Vero cells to maximum cell viability (Fig. 2A,B). SARS-CoV-2 was also susceptible to TP inhibition at the concentrations we tested (Fig. 2C); this was in line with recent reports on the activity of TP against SARS-CoV-2 although our calculated EC_50_ (55.5 µM) is higher than what was reported (2.0412.12 µM) (Yu et al., 2022). Overall, our data suggest that TP can inhibit infection of several coronaviruses *in vitro*.

**Fig. 2.**
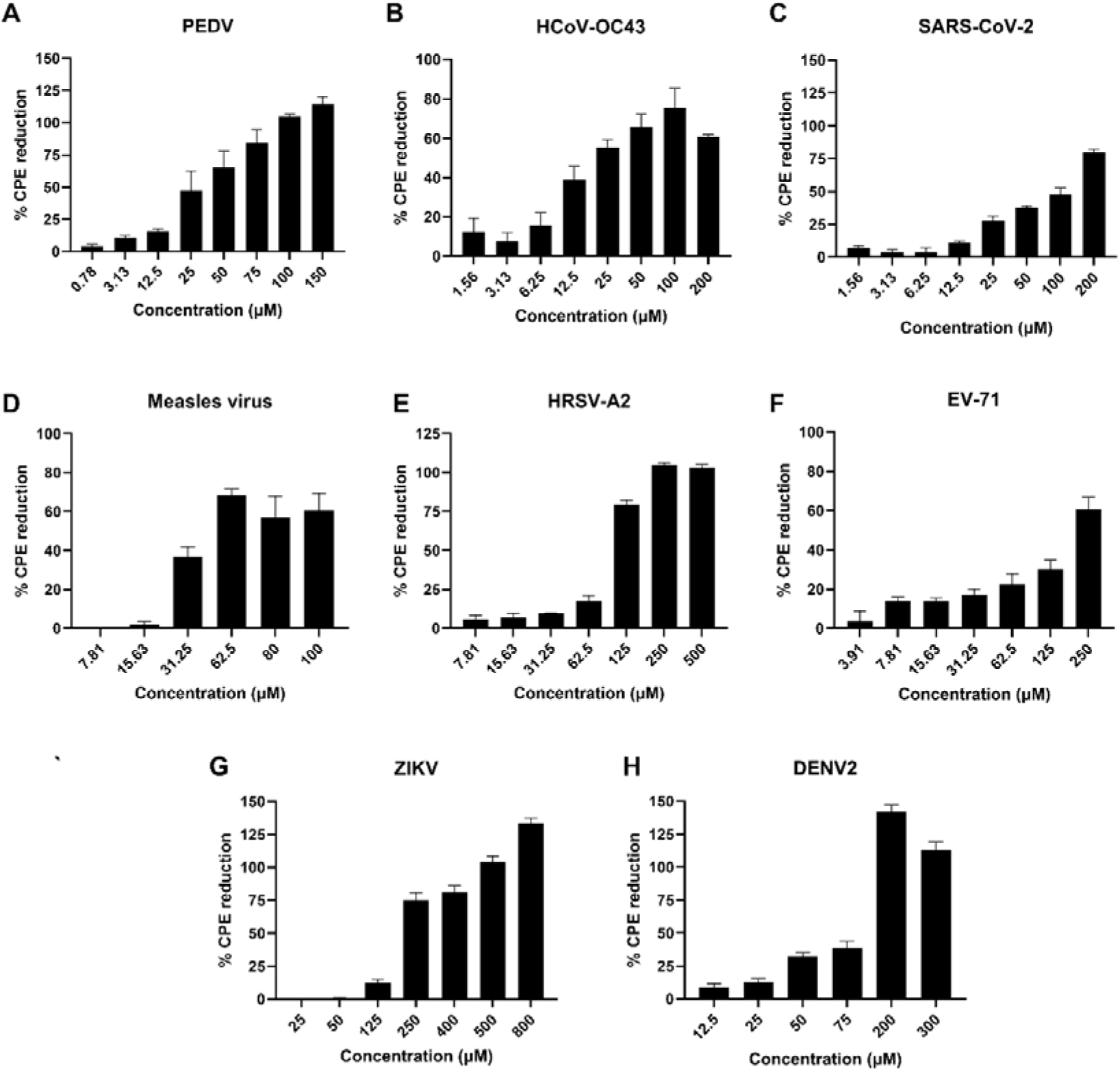
Effects of teicoplanin (TP) on cytopathic effects (CPE) induced by viruses belonging to different virus families. Indicated concentrations of TP were diluted in the assay medium for each virus and used to treat **A,** Vero cells infected with porcine epidemic diarrhea virus (PEDV) at an MOI of 0.001; **B,** Mv1Lu cells infected with 200 TCID_50_/well of human coronavirus OC43 (HCoV-OC43), **C,** Vero E6 cells infected with 100 TCID_50_/well of severe acute respiratory syndrome coronavirus 2 (SARS-CoV-2); **D**, Vero cells infected with 500 TCID_50_/well of measles virus; **E**, HEp-2 cells infected with 70 TCID_50_/well human respiratory syncytial virus A2 (HRSV-A2); **F,** Vero cells infected with 500 TCID_50_/well of enterovirus 71 (EV-71); **G,** Vero cells infected with Zika virus (ZIKV) at an MOI of 0.02; and **H,** Vero E6 cells infected with dengue virus serotype 2 (DENV2) at an MOI of 0.02. Percent cytopathic effect reduction (% CPE reduction) was calculated relative to the viability of the untreated infected and uninfected controls (Ctrl). Data show means ± SEM; n = 3 per TP concentration. Each graph represents one of at least two independent assays

### 3.3 Teicoplanin reduced CPE induced by measles virus and HRSV-A2

Given that TP could inhibit two RNA virus families that cause respiratory infections, we next tested whether the breadth of TP activity could be expanded to other RNA virus families (*Paramyxoviridae* and *Pneumoviridae*) that cause respiratory illness in humans. We chose measles virus as a representative paramyxovirus and HRSV-A2 as a representative pneumovirus. Both viruses were susceptible to inhibition by non-cytotoxic TP concentrations. TP showed complete antiviral activity against HRSV-A2, but it was not able to completely rescue the host cells infected with measles virus to 100% cell viability (Fig. 2D,E). Our results suggest that, in terms of maximal CPE reduction, TP has considerable or moderate ability to inhibit HRSV-A2 or measles virus infection, respectively, further expanding the potential targets of TP antiviral activity.

### 3.4 Teicoplanin reduced CPE induced by enterovirus 71

Thus far, all the RNA viruses we have tested are enveloped RNA viruses. To further expand the breadth of TP targets, we determined the effects of various concentrations on the CPE induced by EV-71 as a representative non-enveloped RNA virus. TP also partially attenuated EV-71 infection in Vero cells (Fig. 2F), suggesting that the effects of TP are not limited to enveloped RNA viruses.

### 3.5 Teicoplanin reduced flavivirus-induced cytopathic effects

Flaviviruses utilize class II fusion proteins, unlike the other enveloped RNA viruses we have tested (influenza viruses, coronaviruses, paramyxovirus, and pneumovirus), which carry class I fusion proteins (Rey and Lok, 2018). TP has been proposed to inhibit cathepsins, especially CatL, that cleave the fusion proteins of SARS-CoV-2 and EBOV (Yu et al., 2022). Since flavivirus E proteins do not require cleavage to facilitate viral entry and host membrane fusion, we presumed that TP might not be able to inhibit flavivirus infection. Upon testing, however, we found that TP could also attenuate CPE induced by both ZIKV and DENV2 in a dose-dependent manner at non-cytotoxic concentrations (Fig. 2G,H). The concentrations needed for complete CPE inhibition of DENV2 were lower than those needed for complete inhibition of ZIKV; however, two different cell lines were used as host for the two viruses, so we cannot directly conclude whether either virus is more susceptible to TP inhibition than the other. Overall, our data show that flaviviruses are also susceptible to TP inhibition.

### 3.6 Teicoplanin did not inhibit human adenovirus 5 infection in vitro

To determine whether TP could inhibit infection of non-enveloped DNA viruses as well, we used human adenovirus 5 (HAd5) as a representative and tested the effects of different TP concentrations against HAd5. TP was unable to rescue HAd5-infected A549 cells, even at the highest concentration (500 mM) we tested (Supplementary Fig. S2).

### 3.7 Addition of TP at the endocytosis stage of PEDV infection reduced PEDV production

To understand the mechanisms underlying the effects of TP on coronaviruses, we first determined the effects of TP treatment on the production of PEDV particles by Vero cells after several cycles of replication. TP reduced PEDV production by ∼1,000-fold, whereas treatment with CQ, a lysomotropic agent that has previously been shown to inhibit SARS-CoV infection *in vitro*, reduced PEDV production by ∼100-fold (Fig. 3A) (Vincent et al., 2005). In a plaque-based assay where we added TP at the virus attachment/binding stage of infection to the host monolayer culture, we found that TP addition did not significantly reduce the production of PEDV virus particles (Fig. 3B). Meanwhile, addition of TP during the endocytosis stage of infection reduced plaque production down to 20%, although, comparably, CQ was still better at impeding virus production when added during the endocytic stage of infection (Fig. 3C). These data suggest that TP does not inhibit binding of PEDV to host cell receptors, instead suppressing viral or host components in the endosomes. These results support previous evidence where TP inhibits endo/lysosomal CatL, leading to reduced levels of activated Spike (S) proteins, thereby impeding fusion of the viral envelope with the host membrane.

**Fig. 3.**
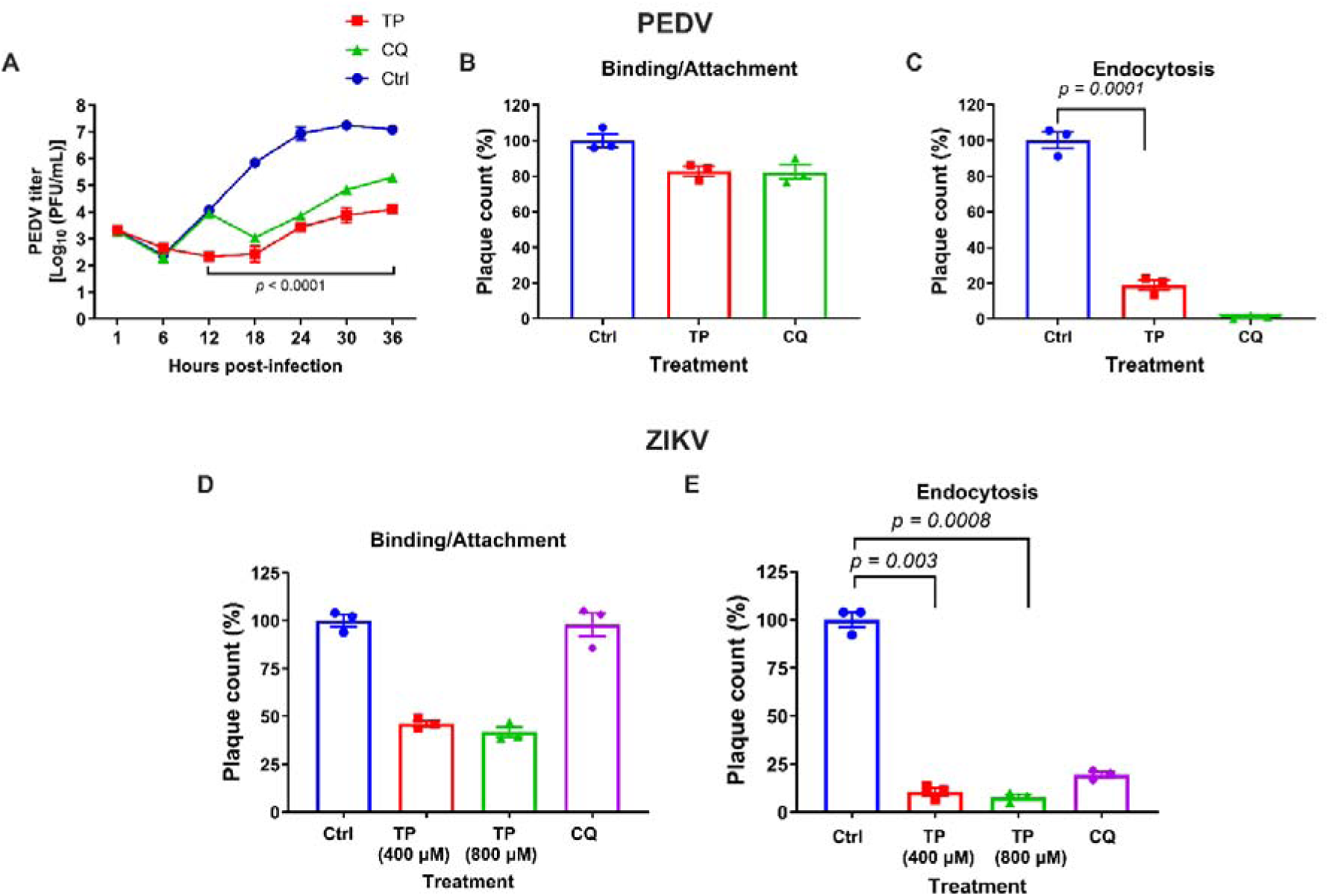
Effects of teicoplanin (TP) treatment on porcine epidemic diarrhea (PEDV) and Zika virus (ZIKV) production *in vitro*. **A**, Vero cells infected with PEDV at an MOI of 0.01 were treated with TP (200 μM) or chloroquine (CQ; 10 μM). Culture supernatants were collected at the indicated timepoints and subjected to plaque assay for PEDV titer determination treatment). **B,** Vero cells were infected with PEDV (200 plaque forming units, PFU) in the presence of 200 μM TP, 10 μM CQ or assay medium (1 mL total volume) for 2 h at 4 °C. The virus + drug mixture was removed, and plaque overlay was added. **C,** Vero cells were infected with 200 PFU of PEDV for 2 h at 4 °C. The inoculum was removed, the cells washed, and 1 mL of 200 μM TP, 10 μM CQ or assay medium was added to the infected cells. The cells were incubated for 2 h at 37 °C, after which, the cells were washed, and plaque overlay was added. **D,** Vero cells were infected with 400 PFU of ZIKV in the presence of 400 μM TP, 800 μM TP, 25 μM CQ or assay medium (1 mL total volume) for 2 h at 4 °C. The virus + drug mixture was removed, and plaque overlay was added. **E,** Vero cells were infected with 400 PFU of ZIKV for 2 h at 4 °C. The inoculum was removed, the cells washed, and 1 mL of 400 μM TP, 800 μM TP, 25 μM CQ or assay medium was added to the infected cells. The cells were incubated for 2 h at 37 °C, after which, the cells were washed, and plaque overlay was added. Plaque count (%) were all calculated relative to the untreated, infected controls. All values are means ± SEM, n = 3 per treatment. All *p-*values are based on Student’s *t-*test; all *p-*values were calculated between TP treatment groups and the untreated infection controls (Ctrl).

### 3.8 TP targets early entry events in the ZIKV replication cycle

In contrast to the effects of TP on PEDV early entry events, we found that TP partially attenuated ZIKV production in a dose-dependent manner when TP was added during the attachment/binding stage of infection, while CQ expectedly did not affect ZIKV production when added at this stage (Fig. 3D). Addition of TP to the ZIKV-infected cells during the endocytosis stage of infection led to an even more dramatic reduction in ZIKV production compared to when TP was added during viral attachment/binding, leading to 90% reduction in plaque formation. CQ, as expected, also inhibited ZIKV endocytosis (Fig. 3E). These results suggest that TP affects early entry events of the ZIKV replication cycle, and its effects are not limited to the endocytosis stage of ZIKV infection.

### 3.9 TP has potential binding pockets in the ZIKV and DENV2 envelope proteins

The class I fusion proteins and other glycoproteins involved in attachment are potential targets for cleavage by cathepsins and are definitive targets of cathepsins in the case of highly pathogenic betacoronaviruses (SARS-CoV, MERS-CoV, and SARS-CoV-2) and filoviruses as shown by previous studies. In contrast, flaviviruses carry a class II fusion protein, the E protein, which does not require proteolytic cleavage to facilitate virus-host fusion; instead, low pH triggers conformational changes to the E proteins (Rey and Lok, 2018). Furthermore, inhibition of furin but not of serine and cysteine proteases has been shown top attenuate ZIKV production *in vitro*, which implies that cathepsins are not critical to ZIKV infection (Owczarek et al., 2019). Despite these, however, ZIKV and DENV2 were susceptible to inhibition by TP. The E protein is critical to entry of flaviviruses into the host. It facilitates attachment to host factors and binding to receptors on the host cell; upon clathrin-dependent endocytosis, the low environmental pH in the endosome triggers reorganization of the E protein, exposing the hydrophobic fusion loop. The fusion loop is then embedded into the host endosomal membrane, and the E proteins assemble into trimers, and succeeding changes to the conformation of the E protein lead to the viral and host membrane fusion, fusion pore formation, and eventual release of the viral genome into the cytoplasm (Stiasny et al., 2023). Any alteration to the flavivirus E protein can therefore disrupt both attachment and fusion stages of flavivirus infection. We thus hypothesized that TP binds the flavivirus E protein, thereby affecting all its functions.

Predicting the potential binding sites of TP on the prefusion form of ZIKV and DENV2 E proteins using PyRx showed that TP may occupy the cavities on either side of the central dimer region of the ZIKV and DENV2 E proteins (Fig. 4). Notably, the top predicted model for the ZIKV E protein with the lowest binding affinity (–8.2 kcal/mol) showed that TP may form hydrogen bonds with ASP99 and GLY103 (Fig. 4B; Supplementary Table S1), which are parts of the fusion loop on one of the E protein chains (chain B). TP was also predicted to form hydrogen bonds with ARG3, SER8, ASN9, GLY29, GLU275, ASP279, and SER286 on one chain (chain A) and with ASP248 on the other chain (chain B). TP was also predicted to have hydrophobic interactions with HIS28 and electrostatic interactions with GLU277 on chain A and electrostatic interactions with ASP248 on chain B. Notably, the predicted TP binding pocket lies on the exposed side of the E protein dimer, allowing TP easier access to the interacting residues. Additionally, because the prefusion conformation of the flavivirus E protein forms homodimers on the surface of the flavivirus particle, TP can potentially bind both pockets on either side of the E protein central dimer region, further impeding movement of the E protein.

**Fig. 4.**
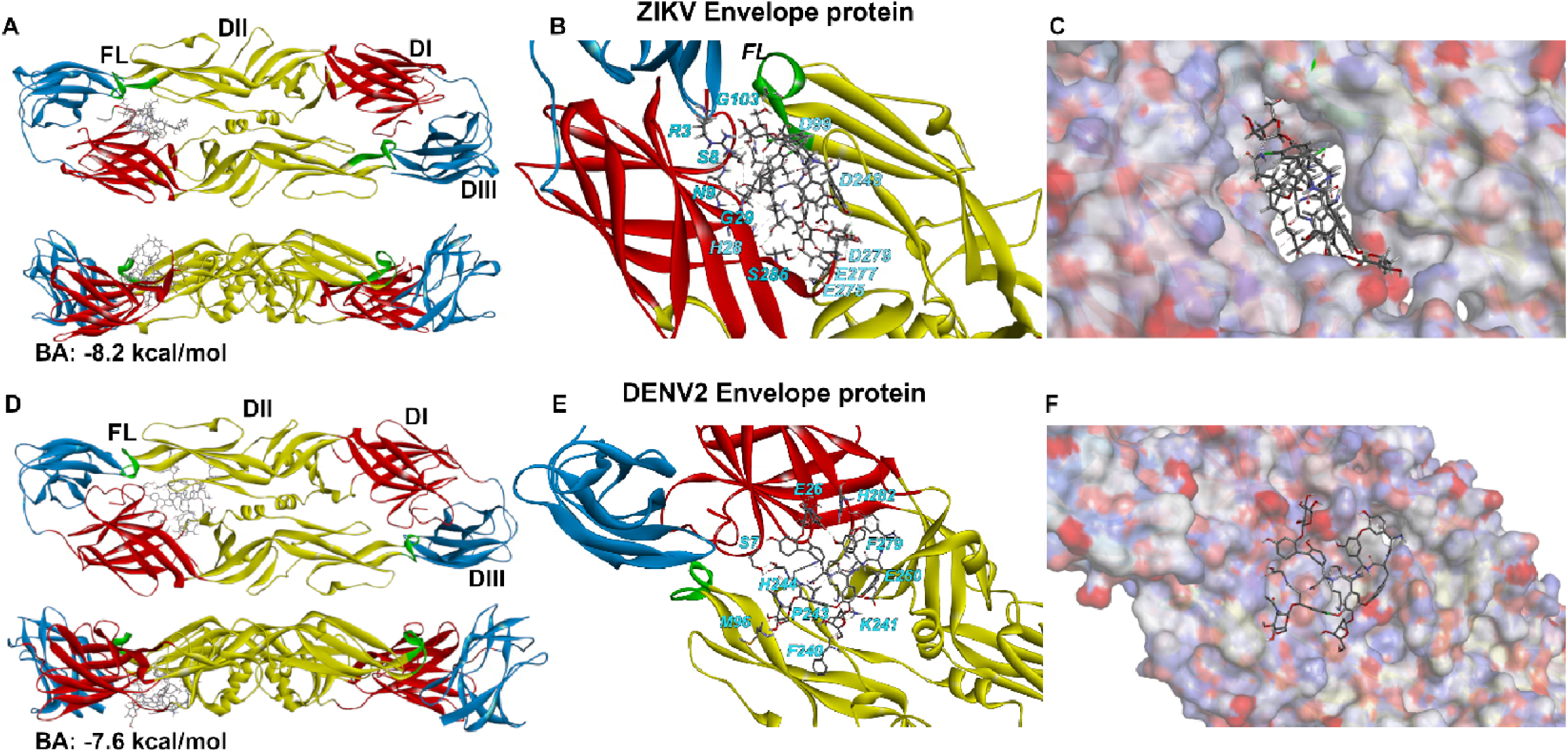
Potential binding pockets of teicoplanin (TP) on the Zika virus (ZIKV) and dengue virus serotype 2 (DENV2) envelope (E) protein. PyRx was to determine the most probable binding sites of TP on the prefusion forms of the **A–C**, ZIKV E protein (PDB ID: 5JHM) and **D–F**, DENV2 E protein ID: 1OKE) using the X-ray crystal structures of the E protein ectodomains. On **A & D**, labeled are the 3 domains of the E proteins: DI (red), DII w), DIII (blue), and the fusion loop (FL; green). The top (top) and the side (bottom) views of the E protein ribbon renderings are shown. For **B & E**, the ues of the ZIKV and DENV2 E proteins that come into contact with TP are shown. **C & F** show the surface charge model of the ZIKV and DENV2 E ins at the potential TP binding pockets, where red regions are negatively charged, and blue regions are positively charged. BA: predicted binding affinity.

Likewise, TP was predicted to bind the cavities near the central dimer region of the prefusion form of the DENV2 E protein (Fig. 4); however, the binding pocket appears to be on the side of the E protein proximal to the viral envelope. TP was predicted to form hydrogen bonds with SER7, and GLU26 on one chain (chain A) and with MET96, LYS241, and PHE240 on the other chain (chain B) of the DENV2 E protein (Fig. 4E; Supplementary Table S2). TP was predicted to have hydrophobic interactions with PHE279 and HIS282 on chain A and HIS244 and PRO243 on chain B of the DENV2 E protein. Meanwhile, the Cl on the structure of TP was predicted to form a halogen interaction with GLU269 on chain A. These interactions could potentially occur on either side of the central dimer region owing to the dimeric structure of the prefusion form of the DENV2 E protein.

## 4 Discussion

Collectively, our results show that TP has a broad range of viral targets (Table 1). This includes enveloped RNA viruses belonging to different virus families: influenza viruses, coronaviruses, flaviviruses, paramyxoviruses, and pneumoviruses. A representative picornavirus (EV-71) was also partially inhibited by TP. However, the only DNA virus we tested (HAd5) was not inhibited by TP treatment. The effects of TP on coronaviruses PEDV and HCoV-OC43 were expected, as a previous study has shown that TP disrupts the entry of MERS-CoV and SARS-CoV spike (S) protein pseudotyped particles and of authentic SARS-CoV-2 particles. In general, the S proteins of alphacoronaviruses and betacoronaviruses require cleavage either on the cell surface or in the acidified endosome, depending on protease availability, which is typically based on host tissue type (Whittaker et al., 2021). In general, wild-type PEDV isolates require the addition of exogenous trypsin to activate the S protein to facilitate propagation in cell culture. However, the PEDV strain that we used (DR13) is trypsin-independent, although trypsin enhances infection (Tan et al., 2022). We did not add trypsin to our cell culture system, and this may have favored the endocytic route, and, in turn cathepsin use, for PEDV S protein activation, allowing TP to fully attenuate PEDV infection in Vero cells. This is further supported by the ability of TP to impede PEDV infection when added during the endocytic stage of infection. Our results corroborate a previous report that the PEDV S protein can be activated by CatL and CatB (Liu et al., 2016). It would be interesting to determine whether TP could still inhibit PEDV infection in the presence of trypsin. Meanwhile, TP was only partially able to attenuate HCoV-OC43 infection in Mv1Lu cells. A previous study has suggested that wild-type HCoV-OC43 uses cell-surface proteases, whereas cell-culture adapted HCoV-OC43 may use the combination of cell-surface proteases and endosomal cathepsins (Shirato et al., 2018). The S protein of the HCoV-OC43 strain we used may be primed through both routes, explaining why the effect of TP was only partial. Meanwhile, the EC_50_ value of TP against SARS-CoV-2 CPE that we calculated was much higher than the previously reported IC_50_ of TP against the Wuhan (2.04 µM) and D614G (2.12 µM) variants of SARS-CoV-2 (Yu et al., 2022). However, our assay system was different from that in the other study, where we used Vero E6 cells and used cell viability as a measure for TP effectivity, while the group in the previous study used hACE2-expressing HEK293T cells and RNA copy numbers as a measure for effectivity. Thus, our results can be taken to corroborate the results of the previous study. Taken together, our results suggest that TP has potential as a pan-coronavirus antiviral candidate, likely through cathepsin inhibition.

The influenza viruses we tested, which include human seasonal and avian subtypes, were all susceptible to TP inhibition. Previous studies have reported that derivatives of TP and glycopeptide antibiotics exhibited different levels of activity against influenza A and B viruses, although a specific target for glycopeptide analogues has not been established (Acharya et al., 2022; Szucs et al., 2018). Additionally, these studies did not test the effects of unmodified TP. In general, activation of the influenza precursor hemagglutinin (HA0) is facilitated by trypsin-type proteases, with HPAI viruses able to utilize furin as well. Although cleavage of HA0 by cathepsins for early entry has not been reported, cathepsin W (CatW) has been demonstrated to be necessary for the fusion of the influenza viral and endosomal membrane fusion (Edinger et al., 2015). CatW did not cleave viral glycoproteins, but the proteolytic activity of cathepsins was still required for the late endosome step, indicating that CatW processes host proteins that promote fusion of the virus with the host. Our results then suggest that CatW inhibition using TP is a feasible pan-influenza antiviral strategy. Time-of-drug-addition assays can be performed to determine whether TP can indeed inhibit influenza virus infection when added before or during the viral and host membrane fusion stage of infection.

Measles virus infection *in vitro* was also partially inhibited by TP treatment. The measles virus precursor fusion glycoprotein (F0) is primarily cleaved by furin in the trans-Golgi network during transport to the extracellular surface following virus assembly (Satoh et al., 2015). Thus far, no study has reported the involvement of cathepsins in the activation of the measles virus F0. However, a recent study has shown that cathepsin inhibition reduced the production of infectious particles of feline morbillivirus (FeMV), which belongs to the same genus as measles virus, and that this inhibition can be attributed to the ability of cathepsins to cleave the FeMV F0 precursor protein (Nambulli et al., 2022). Likewise, Nipah and Hendra viruses that belong to a different genus in the *Paramyxoviridae* family have been shown to depend on CatL instead of furin for F0 cleavage (Pager and Dutch, 2005). It is possible that the measles virus F0 can be cleaved by proteases other than furin, although this hypothesis will have to be verified in future studies. It is also likely that cathepsins act on proviral host factors, which may not be indispensable to the infection process of measles virus as indicated by the partial effects of TP. Additionally, TP may also directly bind and inhibit one of the measles virus proteins, thereby attenuating infection.

Meanwhile, HRSV-A2, which we used as a representative pneumovirus in this study, was also susceptible to inhibition by TP, corroborating the results of a previous study wherein TP inhibited the Long strain of HRSV (Wang et al., 2016). CaTL has been reported to cleave the attachment (G) glycoprotein of HRSV-A2 particles grown in Vero cells but not those grown in HeLa cells, which may be due to differences in the glycosylation patterns on the resulting viruses (Corry et al., 2016). This cleavage of the G protein by CatL is proposed to occur during endocytic recycling of the G protein and results in HRSV particles with shorter G proteins, hampering its ability to attach to heparan sulfate on the host cell. Since we used HEp-2 cells, which are derived from HeLa cells, for virus production, there is low likelihood that the HRSV-A2 G proteins generated in our culture system had the glycosylation patterns that make the G protein amenable to CatL cleavage. While it is still likely that CatL could cleave the G protein of the HRSV-A2 we produced, CatL cleavage should reduce the infectivity of HRSV, and CatL inhibition should in turn promote HRSV infectivity in host cells. This was the opposite of what we observed, suggesting that: a) CatL cleaves another HRSV protein, b) CatL plays a role in proviral pathway that becomes downregulated by TP treatment, or c) TP directly interacts with and inhibits the function of a viral protein. The mechanisms underlying the susceptibility of HRSV to TP treatment should be explored in future studies.

In contrast to the results of a previous study, we found TP was partially able to inhibit infection of EV-71 in host cells at non-cytotoxic concentrations (Wang et al., 2016). These different results may be due to the different cell lines used, where we used Vero cells and the other group used RD cells as host for EV-71. Additionally, the TP concentrations we tested were higher, as Vero cells were more tolerant of TP than RD cells, based on the effects of TP on the viability of both cell lines as we present here and as presented in the previous study. Regardless, it appears that EV-71 has moderate susceptibility to TP treatment. CatB has been linked to viral myocarditis in mice infected with the related coxsackievirus B3; however, the study presented an inflammatory role for CatB, which may not be applicable to our study system (Wang et al., 2018). It is still unclear whether cathepsins are involved specifically in the replication cycle of EV-71 and other picornaviruses. Picornavirus genomes encode a 3 chymotrypsin-like cysteine protease (3CLpro). Considering that cathepsins are also cysteine proteases, TP may also be able to inhibit the picornavirus 3CLpro, although this is so far an unexplored property of TP. A study in 2011 has shown that the poliovirus 3CLpro was susceptible to tellurium-based compounds that were also capable of inhibiting CatB, CatL, CatS, and CatK (Gouvea et al., 2011). It is possible that the picornavirus 3CLpro shares proteolytic active sites with cathepsins that makes it susceptible to inhibition by TP and other cysteine protease inhibitors.

While TP was unable to impede PEDV infection when added at the attachment/binding stage but inhibited PEDV infection when added at the endocytosis stage, TP was partially able to inhibit ZIKV production when added to the host during the attachment/binding stage and completely inhibited ZIKV production when added at the endocytosis stage of infection. Our molecular docking simulations suggest that TP binds pockets near the central dimer regions of the E proteins of ZIKV and DENV, with TP potentially binding the fusion loop of the ZIKV protein. Addition of TP at the early entry events of flavivirus infection may lock the E protein of the virus to the dimeric state and, especially in the case of ZIKV, block the interaction of the ZIKV E protein fusion loop with the host membrane, thus preventing fusion. The differences between the effects of TP when added during the attachment and the endocytic stages of infection may be attributed to the different states of the E protein, wherein TP may be more likely to interact with the endocytic state of the E protein than to the prefusion form, perhaps with more exposed TP binding sites when the E protein is in the low-pH state. Alternatively, the changes that TP binding induces on the E protein conformation may not be critical to E protein binding to the receptor but may be indispensable to the fusion machinery, and that the TP-bound E proteins during the attachment stage carry over to the fusion stage, thereby impeding virus-host membrane fusion. Previous studies have also associated NS1 with the upregulation of CatL in endothelial cells upon infection with various flaviviruses. However, this effect was specific to the tissues associated with the pathogenesis of each virus (i.e., human umbilical vein and human brain microvascular endothelial cells for ZIKV; human brain microvascular endothelial cells for WNV and JEV; human liver sinusoidal endothelial cells for YFV; and various cell types for DENV) and would therefore not be evident on Vero cells (Puerta-Guardo et al., 2019). The JEV capsid protein has also been reported to be susceptible to processing by CatL (Mori et al., 2007). However, addition of a CatL inhibitor attenuated JEV virus production to virus-infected mouse macrophage and neuroblastoma cells but not in virus-infected Vero cells, indicating tissue-specificity as well. Furthermore, the potential interactions of the flaviviral NS1 or C protein do not explain the attenuation of ZIKV production when TP was added during the attachment/binding stage of infection. A study has also reported that a furin inhibitor but neither serine nor cysteine protease inhibitors reduced virus production in ZIKV-infected cells, which further supports the idea that cathepsins are not required by ZIKV for infection (Owczarek et al., 2019). Thus, at least in our assay system, the most probable target of TP would be the E protein. Supporting this, an aglycone derivative of TP has been reported to inhibit flavivirus infection, including that of DENV by impeding binding of the DENV particles to host cells (De Burghgraeve et al., 2012).

TP is primarily used as an antibiotic for most Gram-positive bacteria. Bacterial co-infections and secondary co-infections are common in late stages of viral respiratory tract infections. Frequency of bacterial co-infections with influenza virus infection is particularly high, with estimates within 20.31123% regardless of age group, and increases the risk of death by around 3.4 times among hospitalized patients relative to influenza virus infection alone (Arranz-Herrero et al., 2023; Qiao et al., 2023). Studies report that the most common co-infecting bacteria during influenza virus infections are *S. aureus* and *S. pneumoniae* (Oliva and Terrier, 2021; Wong et al., 2013). Co-infections of RSV and bacteria (*S. aureus, S. pneumoniae, and Haemophilus influenzae*) have also been reported (17.5 LJ 44%) among hospitalized patients and have been correlated with increased severity compared to lone RSV infection (Lee et al., 2013; Oliva and Terrier, 2021; Pacheco et al., 2021). On the other hand, estimates on the rates of bacterial co-infections with SARS-CoV-2 have been lower than those with influenza virus and RSV; however, bacterial co-infections and superinfections with SARS-CoV-2 have similarly been associated with worse outcomes compared with SARS-CoV-2 infection alone (Garcia-Vidal et al., 2021; Oliva and Terrier, 2021). Measles virus infection can suppress immune responses and damage the epithelium, making infected persons more susceptible to bacterial infections that have been associated with complications of measles, including pneumonia, diarrhea, and otitis media (Rota et al., 2016). Antibiotic use in addition to antiviral use is recommended for confirmed cases of bacterial pneumonia following influenza virus infection (Fiore et al., 2011). TP, especially in cases of MRSA co-infections and superinfections, may then be considered a good option for treatment over other antibiotics that do not have antiviral effects. However, antibiotic resistance is a growing problem worldwide, making TP a double-edged sword as an antiviral candidate. On one hand, it may be able to prevent or minimize the effects of bacterial co-infections and superinfections when used as an antiviral; on the other hand, inappropriate usage may lead to the emergence of antibiotic-resistant bacteria. TP can instead serve as a parent compound for the development of non-antibiotic antivirals.

TP is a large, complex semisynthetic glycopeptide consisting of five main structures. It potentially contains multiple pharmacophores with antiviral effects, which may account for the range of its viral targets. Other glycopeptide antibiotics probably share these pharmacophores as they have also been reported to exhibit antiviral effects at varying degrees (Acharya et al., 2022). Indeed, studies on TP derivatives suggest that the different modifications lead to different levels of activity against influenza virus infection (Acharya et al., 2022; Szucs et al., 2018). This is also seen in studies on the effects of the derivatives of TP and other glycopeptides on hepatitis C virus (HCV) infection, where one derivative seemed to inhibit viral genome replication, which is in contrast to the inhibitory effects of glycopeptide antibiotics on the entry of other viruses (Obeid et al., 2011). Identifying the specific pharmacophores with different antiviral mechanisms, for example one pharmacophore that inhibits cathepsins and another that binds flavivirus envelope proteins, and developing them into compounds with more potent and more specific antiviral activity without antibacterial effects may be a better strategy for taking advantage of the broad antiviral range of TP. Additionally, the effects of TP and other glycopeptide antibiotics on viral infection in human cases should be evaluated. In line with this, TP treatment on a patient with chronic hepatitis C was reported to have reduced HCV RNA copy numbers over the course of TP treatment following surgery, and the HCV RNA copy numbers elevated after cessation of TP treatment (Maieron and Kerschner, 2012). Further studies should be performed to determine whether this observation can be replicated.

## 5 Conclusions

Here, we show that TP inhibited several RNA viruses across different families with varying levels of effectivity but did not inhibit a representative non-enveloped DNA virus. Remarkably, TP inhibited all the representative RNA viruses we tested with or without the viral lipid envelope. Furthermore, TP inhibited RNA viruses with either class I or class II fusion proteins. A closer look at the effects of TP on early entry events of PEDV suggested that TP inhibits PEDV endocytosis but not PEDV attachment to host cells. Meanwhile, TP was able to completely inhibit ZIKV endocytosis and was able to partially inhibit ZIKV attachment to host cells. This might point to different mechanisms of action of TP against coronavirus and flavivirus infection, which may allude to differences in viral fusion mechanisms as coronaviruses bear class I fusion proteins, while flaviviruses bear class II fusion proteins. TP may be able to inhibit the entry of viruses with class I fusion proteins endocytosis through inhibition of cathepsins required to cleave and activate class I fusion proteins. Meanwhile, TP may be able to inhibit flavivirus entry by binding and inhibiting the flavivirus envelope protein. TP may also be able to inhibit a step of endocytosis by inhibiting yet unknown mechanisms in the infection cycle of flaviviruses. Whether TP can inhibit other RNA viruses with class II fusion proteins should also be tested on Alphaviruses. All in all, our study shows that TP consists of pharmacophores that allow it to inhibit a broad range of RNA viruses. Further studies should be performed to determine the specific targets of TP and the corresponding pharmacophore. Notably, TP inhibits most viruses that cause severe respiratory illness in humans that are also linked to secondary bacterial infection. Whether TP can be utilized as treatment for these respiratory illnesses in regimens that minimize the risk of bacterial co-infections without contributing to the growing problem of antibiotic resistance should be evaluated. However, to minimize the potential emergence of antibiotic-resistant bacterial strains, the pharmacophore of TP with broad antiviral activity should be identified and developed into a safe, non-antibacterial product for clinical use.

## Ethics statement

No animals were used in this study. No patients were involved in this study.

## Availability of data and materials

All data generated during the current study are included in the manuscript.

## CRediT authorship contribution statement

**Erica Españo:** Writing - Original draft, Methodology, Validation, Formal analysis, Investigation, Data curation, Visualization. **Jiyeon Kim:** Writing - Original draft, Methodology, Validation, Formal analysis, Investigation, Data curation. **Seong Ok Park:** Methodology, Investigation. **Bill Thaddeus Padasas:** Investigation. **Sang-Hyun Kim:** Investigation. **Ju-Ho Son:** Investigation. **Jihee Oh:** Investigation. **E-Eum Woo:** Investigation. **Young-Ran Lee:** Writing - review & editing, Resources. **Soon Young Hwang:** Writing - review & editing, Conceptualization, Resources, Funding acquisition. **Seong Kug Eo:** Writing - review & editing, Validation, Resources, Supervision. **Jeong-Ki Kim:** Writing - review & editing, Conceptualization, Validation, Resources, Supervision, Project administration, Funding acquisition.

## References

1. Acharya, Y., Bhattacharyya, S., Dhanda, G. & Haldar, J., 2022. Emerging roles of glycopeptide antibiotics: Moving beyond Gram-positive bacteria. ACS Infect. Dis. 8, 1–28. 10.1021/acsinfecdis.1c00367.00

2. Arranz-Herrero, J., Presa, J., Rius-Rocabert, S., Utrero-Rico, A., Arranz-Arija, J.A., Lalueza, A., Escribese, M.M., Ochando, J., Soriano, V. & Nistal-Villan, E., 2023. Determinants of poor clinical outcome in patients with influenza pneumonia: A systematic review and meta-analysis. Int. J. Infect. Dis. 131, 173–179. 10.1016/j.ijid.2023.04.003.

3. Balzarini, J., Pannecouque, C., De Clereq, E., Pavlov, A.Y., Printsevskaya, S.S., Miroshnikova, O.V., Reznikova, M.I. & Preobrazhenskaya, M.N., 2003. Antiretroviral activity of semisynthetic derivatives of glycopeptide antibiotics. J. Med. Chem. 46, 2755–2764. 10.1021/jm0300882.

4. Cerqueira, C., Ventayol, P.S., Vogeley, C. & Schelhaas, M., 2015. Kallikrein-8 proteolytically processes human papillomaviruses in the extracellular space to facilitate entry into host cells. J. Virol. 89, 7038–7052. 10.1128/Jvi.00234-15.

5. Chandran, K., Sullivan, N.J., Felbor, U., Whelan, S.P. & Cunningham, J.M., 2005. Endosomal proteolysis of the Ebola virus glycoprotein is necessary for infection. Science 308, 1643–1645. 10.1126/science.1110656.

6. Corry, J., Johnson, S.M., Cornwell, J. & Peeples, M.E., 2016. Preventing cleavage of the respiratory syncytial virus attachment protein in Vero cells rescues the infectivity of progeny virus for primary human airway cultures. J. Virol. 90, 1311–1320. 10.1128/JVI.02351-15.

7. Dallakyan, S. & Olson, A.J., 2015. Small-molecule library screening by docking with PyRx. Methods Mol. Biol. 1263, 243–250. 10.1007/978-1-4939-2269-7_19.

8. De Burghgraeve, T., Kaptein, S.J.F., Ayala-Nunez, N.V., Mondotte, J.A., Pastorino, B., Printsevskaya, S.S., de Lamballerie, X., Jacobs, M., Preobrazhenskaya, M., Gamarnik, A.V., Smit, J.M. & Neyts, J., 2012. An analogue of the antibiotic teicoplanin prevents flavivirus entry in vitro. PLoS One 7, e37244. 10.1371/journal.pone.0037244.

9. Eastman, R.T., Roth, J.S., Brimacombe, K.R., Simeonov, A., Shen, M., Patnaik, S. & Hall, M.D., 2020. Remdesivir: A review of its discovery and development leading to emergency use authorization for treatment of COVID-19. ACS Central. Sci. 6, 672–683. 10.1021/acscentsci.0c00489.

10. Ebert, D.H., Deussing, J., Peters, C. & Dermody, T.S., 2002. Cathepsin L and cathepsin B mediate reovirus disassembly in murine fibroblast cells. J. Biol. Chem. 277, 24609–24617. 10.1074/jbc.M201107200.

11. Edinger, T.O., Pohl, M.O., Yángüez, E. & Stertz, S., 2015. Cathepsin W is required for escape of influenza A virus from late endosomes. mBio 6, e00297. 10.1128/mbio.00297-15.

12. Fiore, A.E., Fry, A., Shay, D., Gubareva, L., Bresee, J.S., Uyeki, T.M., Centers for Disease, C. & Prevention, 2011. Antiviral agents for the treatment and chemoprophylaxis of influenza --- recommendations of the Advisory Committee on Immunization Practices (ACIP). MMWR Recomm. Rep. 60, 1–24.

13. Garcia-Vidal, C., Sanjuan, G., Moreno-García, E., Puerta-Alcalde, P., Garcia-Pouton, N., Chumbita, M., Fernandez-Pittol, M., Pitart, C., Inciarte, A., Bodro, M., Morata, L., Ambrosioni, J., Grafia, I., Meira, F., Macaya, I., Cardozo, C., Casals, C., Tellez, A., Castro, P., Marco, F., García, F., Mensa, J., Martínez, J.A., Soriano, A. & Grp, C.-R., 2021. Incidence of co-infections and superinfections in hospitalized patients with COVID-19: a retrospective cohort study. Clin. Microbiol. Infect. 27, 83–88. 10.1016/j.cmi.2020.07.041.

14. Gouvea, I.E., Santos, J.A.N., Burlandy, F.M., Tersariol, I.L.S., da Silva, E.E., Juliano, M.A., Juliano, L. & Cunha, R.L.O.R., 2011. Poliovirus 3C proteinase inhibition by organotelluranes. Biol. Chem. 392, 587–591. 10.1515/Bc.2011.059.

15. Imran, M., Arora, M.K., Asdaq, S.M.B., Khan, S.A., Alaqel, S.I., Alshammari, M.K., Alshehri, M.M., Alshrari, A.S., Ali, A.M., Al-shammeri, A.M., Alhazmi, B.D., Harshan, A.A., Alam, M.T. & Abida, 2021. Discovery, development, and patent trends on molnupiravir: A prospective oral treatment for COVID-19. Molecules 26, 5795. 10.3390/molecules26195795.

16. Lee, N., Lui, G.C.Y., Wong, K.T., Li, T.C.M., Tse, E.C.M., Chan, J.Y.C., Yu, J., Wong, S.S.M., Choi, K.W., Wong, R.Y.K., Ngai, K.L.K., Hui, D.S.C. & Chan, P.K.S., 2013. High morbidity and mortality in adults hospitalized for respiratory syncytial virus infections. Clin. Infect. Dis. 57, 1069–1077. 10.1093/cid/cit471.

17. Liu, C., Ma, Y.M., Yang, Y., Zheng, Y., Shang, J., Zhou, Y.S., Jiang, S.B., Du, L.Y., Li, J.R. & Li, F., 2016. Cell entry of porcine epidemic diarrhea coronavirus is activated by lysosomal proteases. J. Biol. Chem. 291, 24779–24786. 10.1074/jbc.M116.740746.

18. Maieron, A. & Kerschner, H., 2012. Teicoplanin therapy leading to a significant decrease in viral load in a patient with chronic hepatitis C. J. Antimicrob. Chemother. 67, 2537–2538. 10.1093/jac/dks217.

19. Majerová, T. & Konvalinka, J., 2022. Viral proteases as therapeutic targets. Mol. Aspects Med. 88. 10.1016/j.mam.2022.101159.

20. Mercorelli, B., Palù, G. & Loregian, A., 2018. Drug repurposing for viral infectious diseases: How far are we? Trends Microbiol. 26, 865–876. 10.1016/j.tim.2018.04.004.

21. Mori, Y., Yamashita, T., Tanaka, Y., Tsuda, Y., Abe, T., Moriishi, K. & Matsuura, Y., 2007. Processing of capsid protein by cathepsin L plays a crucial role in replication of Japanese encephalitis virus in neural and macrophage cells. J. Virol. 81, 8477–8487. 10.1128/JVI.00477-07.

22. Nam, J.H., Kim, E.H., Song, D., Choi, Y.K., Kim, J.K. & Poo, H., 2011. Emergence of mammalian species-infectious and -pathogenic avian influenza H6N5 virus with no evidence of adaptation. J. Virol. 85, 13271–13277. 10.1128/Jvi.05038-11.

23. Nam, J.H., Shim, S.M., Song, E.J., Espano, E., Jeong, D.G., Song, D. & Kim, J.K., 2017. Rapid virulence shift of an H5N2 avian influenza virus during a single passage in mice. Arch. Virol. 162, 3017–3024. 10.1007/s00705-017-3451-9.

24. Nambulli, S., Rennick, L.J., Acciardo, A.S., Tilston-Lunel, N.L., Ho, G., Crossland, N.A., Hardcastle, K., Nieto, B., Bainbridge, G., Williams, T., Sharp, C.R. & Duprex, W.P., 2022. FeMV is a cathepsin-dependent unique morbillivirus infecting the kidneys of domestic cats. Proc. Natl. Acad. Sci. USA 119, e2209405119. 10.1073/pnas.2209405119.

25. Nyström, K., Waldenström, J., Tang, K.W. & Lagging, M., 2019. Ribavirin: pharmacology, multiple modes of action and possible future perspectives. Future Virol. 14, 153–160. 10.2217/fvl-2018-0166.

26. Obeid, S., Printsevskaya, S.S., Olsufyeva, E.N., Dallmeier, K., Durantel, D., Zoulim, F., Preobrazhenskaya, M.N., Neyts, J. & Paeshuyse, J., 2011. Inhibition of hepatitis C virus replication by semi-synthetic derivatives of glycopeptide antibiotics. J. Antimicrob. Chemother. 66, 1287–1294. 10.1093/jac/dkr104.

27. Oliva, J. & Terrier, O., 2021. Viral and bacterial co-infections in the lungs: dangerous liaisons. Viruses 13, 1725. 10.3390/v13091725

28. Owczarek, K., Chykunova, Y., Jassoy, C., Maksym, B., Rajfur, Z. & Pyrc, K., 2019. Zika virus: mapping and reprogramming the entry. Cell Commun. Signal 17, 41. 10.1186/s12964-019-0349-z.

29. Pacheco, G.A., Gálvez, N.M.S., Soto, J.A., Andrade, C.A. & Kalergis, A.M., 2021. Bacterial and viral coinfections with the human respiratory syncytial virus. Microorganisms 9, 1293. 10.3390/microorganisms9061293.

30. Pager, C.T. & Dutch, R.E., 2005. Cathepsin L is involved in proteolytic processing of the Hendra virus fusion protein. J. Virol. 79, 12714–12720. 10.1128/JVI.79.20.12714-12720.2005.

31. Pierson, T.C. & Diamond, M.S., 2012. Degrees of maturity: the complex structure and biology of flaviviruses. Curr. Opin. Virol. 2, 168–175. 10.1016/j.coviro.2012.02.011.

32. Puerta-Guardo, H., Glasner, D.R., Espinosa, D.A., Biering, S.B., Patana, M., Ratnasiri, K., Wang, C.L., Beatty, P.R. & Harris, E., 2019. Flavivirus NS1 triggers tissue-specific vascular endothelial dysfunction reflecting disease tropism. Cell Rep. 26, 1598–1613. 10.1016/j.celrep.2019.01.036.

33. Qiao, M.L., Moyes, G., Zhu, F.Y., Li, Y. & Wang, X., 2023. The prevalence of influenza bacterial co-infection and its role in disease severity: A systematic review and meta-analysis. J. Glob. Health 13, 04063. 10.7189/jogh.13.04063.

34. Rey, F.A. & Lok, S.M., 2018. Common features of enveloped viruses and implications for immunogen design for next-generation vaccines. Cell 172, 1319–1334. 10.1016/j.cell.2018.02.054.

35. Rota, P.A., Moss, W.J., Takeda, M., de Swart, R.L., Thompson, K.M. & Goodson, J.L., 2016. Measles. Nat. Rev. Dis. Primers 2, 16049. 10.1038/nrdp.2016.49.

36. Satoh, Y., Hirose, M., Shogaki, H., Wakimoto, H., Kitagawa, Y., Gotoh, B., Takahashi, K. & Itoh, M., 2015. Intramolecular complementation of measles virus fusion protein stability confers cell-cell fusion activity at 37 °C. FEBS Lett. 589, 152–158. 10.1016/j.febslet.2014.11.040.

37. Scarcella, M., d’Angelo, D., Ciampa, M., Tafuri, S., Avallone, L., Pavone, L.M. & De Pasquale, V., 2022. The key role of lysosomal protease cathepsins in viral infections. Int. J. Mol. Sci. 23, 9089. 10.3390/ijms23169089.

38. Shirato, K., Kawase, M. & Matsuyama, S., 2018. Wild-type human coronaviruses prefer cell-surface TMPRSS2 to endosomal cathepsins for cell entry. Virology 517, 9–15. 10.1016/j.virol.2017.11.012.

39. Skidmore, A.M. & Bradfute, S.B., 2023. The life cycle of the alphaviruses: From an antiviral perspective. Antiviral Res. 209, 105476. 10.1016/j.antiviral.2022.105476.

40. Song, D.S., Yang, J.S., Oh, J.S., Han, J.H. & Park, B.K., 2003. Differentiation of a Vero cell adapted porcine epidemic diarrhea virus from Korean field strains by restriction fragment length polymorphism analysis of ORF 3. Vaccine 21, 1833–1842. 10.1016/S0264-410x(03)00027-6.

41. Song, E.J., Espano, E., Shim, S.M., Nam, J.H., Kim, J., Lee, K., Park, S.K., Lee, C.K. & Kim, J.K., 2021. Inhibitory effects of aprotinin on influenza A and B viruses *in vitro* and *in vivo*. Sci. Rep. 11, 9427. 10.1038/s41598-021-88886-1.

42. Stiasny, K., Medits, I., Rossbacher, L. & Heinz, F.X., 2023. Impact of structural dynamics on biological functions of flaviviruses. FEBS J. 290, 1973–1985. 10.1111/febs.16419.

43. Svetitsky, S., Leibovici, L. & Paul, M., 2009. Comparative efficacy and safety of vancomycin versus teicoplanin: Systematic review and meta-analysis. Antimicrob. Agents Chemother. 53, 4069–4079. 10.1128/Aac.00341-09.

44. Szucs, Z., Kelemen, V., Thai, S.L., Csávás, M., Roth, E., Batta, G., Stevaert, A., Vanderlinden, E., Naesens, L., Herczegh, P. & Borbás, A., 2018. Structure-activity relationship studies of lipophilic teicoplanin pseudoaglycon derivatives as new anti-influenza virus agents. Eur. J. Med. Chem. 157, 1017–1030. 10.1016/j.ejmech.2018.08.058.

45. Tan, Y.B., Sun, L.M., Wang, G., Shi, Y.J., Dong, W.Y., Fu, Y.N., Fu, Z., Chen, H.C. & Peng, G.Q., 2022. Trypsin-enhanced infection with porcine epidemic diarrhea virus is determined by the S2 subunit of the spike glycoprotein. J. Virol. 95, e02453–20. 10.1128/jvi.02453-20.

46. Unal, M.A., Bitirim, C.V., Summak, G.Y., Bereketoglu, S., Zeytin, I.C., Besbinar, O., Gurcan, C., Aydos, D., Goksoy, E., Kocakaya, E., Eran, Z., Murat, M., Demir, N., Ozer, Z.B.A., Somers, J., Demir, E., Nazir, H., Ozkan, S.A., Ozkul, A., Azap, A., Yilmazer, A. & Akcali, K.C., 2021. Ribavirin shows antiviral activity against SARS-CoV-2 and downregulates the activity of TMPRSS2 and the expression of ACE2 in vitro. Can. J. Physiol. Pharmacol. 99, 449–460. 10.1139/cjpp-2020-0734.

47. Vincent, M.J., Bergeron, E., Benjannet, S., Erickson, B.R., Rollin, P.E., Ksiazek, T.G., Seidah, N.G. & Nichol, S.T., 2005. Chloroquine is a potent inhibitor of SARS coronavirus infection and spread. Virol. J. 2, 69. 10.1186/1743-422x-2-69.

48. Wang, Y.P., Jia, L.L., Shen, J., Wang, Y.D., Fu, Z.R., Su, S.A., Cai, Z.J., Wang, J.A. & Xiang, M.X., 2018. Cathepsin B aggravates coxsackievirus B3-induced myocarditis through activating the inflammasome and promoting pyroptosis. PLoS Pathog. 14, e1006872. 10.1371/journal.ppat.1006872.

49. Wang, Y.Z., Cui, R., Li, G.M., Gao, Q.Q., Yuan, S.L., Altmeyer, R. & Zou, G., 2016. Teicoplanin inhibits Ebola pseudovirus infection in cell culture. Antiviral Res. 125, 1–7. 10.1016/j.antiviral.2015.11.003.

50. Whittaker, G.R., Daniel, S. & Millet, J.K., 2021. Coronavirus entry: how we arrived at SARS-CoV-2. Curr. Opin. Virol. 47, 113–120. 10.1016/j.coviro.2021.02.006.

51. Wong, K.K., Jain, S., Blanton, L., Dhara, R., Brammer, L., Fry, A.M. & Finelli, L., 2013. Influenza-associated pediatric deaths in the United States, 2004-2012. Pediatrics 132, 796–804. 10.1542/peds.2013-1493.

52. Yadati, T., Houben, T., Bitorina, A. & Shiri-Sverdlov, R., 2020. The ins and outs of cathepsins: Physiological function and role in disease management. Cells 9, 1679. 10.3390/cells9071679

53. Yu, F., Pan, T., Huang, F., Ying, R.S., Liu, J., Fan, H.M., Zhang, J.S., Liu, W.W., Lin, Y.T., Yuan, Y.C., Yang, T., Li, R., Zhang, X., Lv, X., Chen, Q.Y., Liang, A.Q., Zou, F., Liu, B.F., Hu, F.Y., Tang, X.P., Li, L.H., Deng, K., He, X., Zhang, H., Zhang, Y.W. & Ma, X.C., 2022. Glycopeptide antibiotic teicoplanin inhibits cell entry of SARS-CoV-2 by suppressing the proteolytic activity of cathepsin L. Front. Microbiol. 13, 884034. 10.3389/fmicb.2022.884034.

54. Zhou, N., Pan, T., Zhang, J.S., Li, Q.W., Zhang, X., Bai, C., Huang, F., Peng, T., Zhang, J.H., Liu, C., Tao, L. & Zhang, H., 2016. Glycopeptide antibiotics potently inhibit cathepsin L in the late endosome/lysosome and block the entry of Ebola virus, Middle East respiratory syndrome coronavirus (MERS-CoV), and severe acute respiratory syndrome coronavirus (SARS-CoV). J. Biol. Chem. 291, 9218–9232. 10.1074/jbc.M116.716100.

