## Supplemental Figures for "Teicoplanin attenuates RNA virus infection *in vitro*"

Erica Españo^a,1^, Jiyeon Kim^a,1^, Seong Ok Park^b^, Bill Thaddeus Padasas^a^, Sang-Hyun Kim^a^, Ju-Ho Son^a^, Jihee Oh^a^, E-Eum Woo^c^, Young-Ran Lee^d^, Soon Young Hwang^c,***^, Seong Kug Eo^b,**^, Jeong-Ki Kim^a,*^

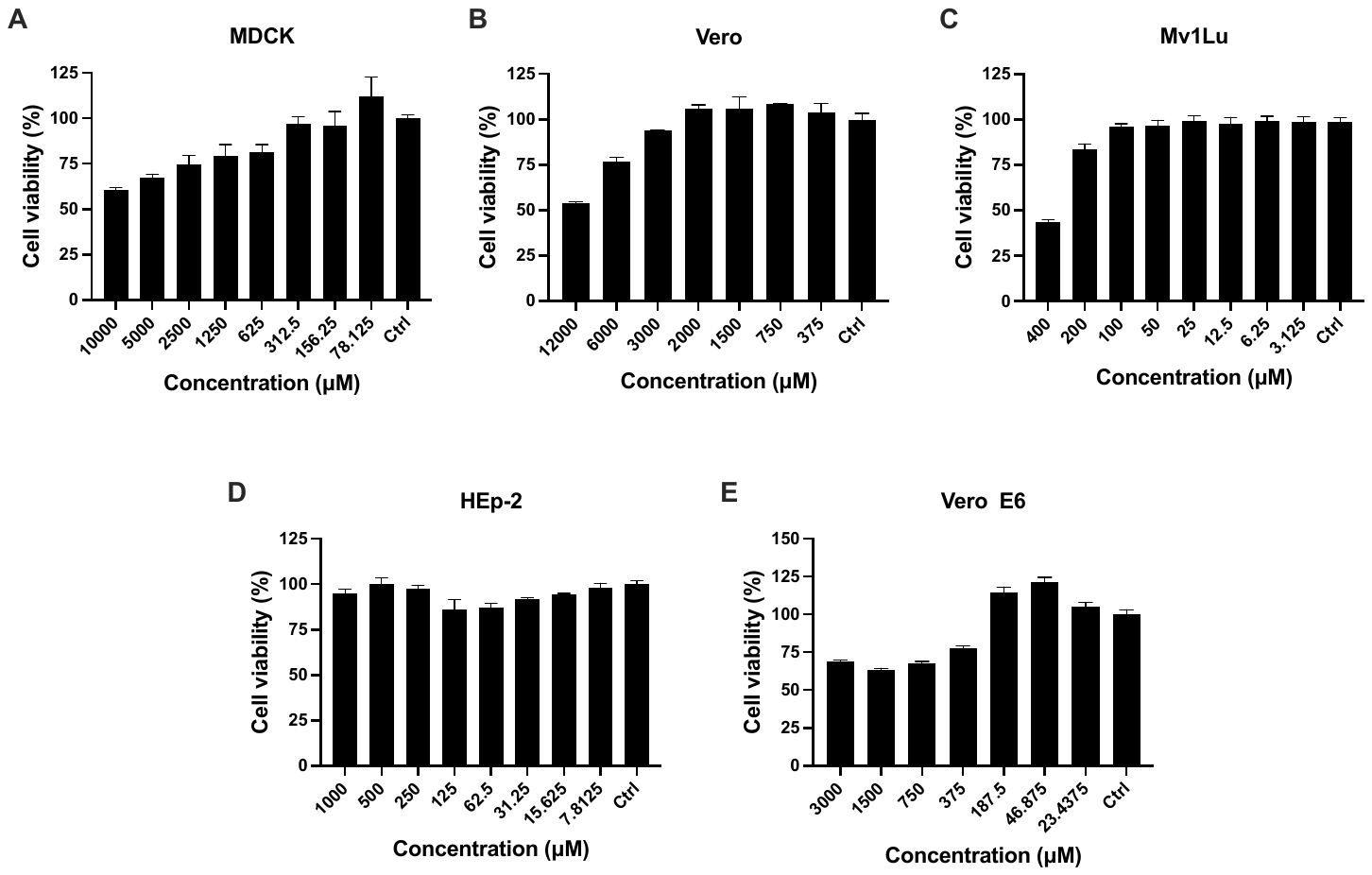

**Supplementary Fig. S1. Effects of teicoplanin (TP) on the viability of the various cell lines that were used as viral hosts for this study.** The following cells were seeded into 96-well plates and then treated with the indicated concentrations of TP to determine the cytotoxicity of TP on each cell line: **A,** MDCK; **B,** Vero; **C,** MV1Lu; **D,** HEp-2; **E,** Vero E6. After the incubation periods corresponding to viral infection, cell viability was determined using the EZ-Cytox reagent. All % cell viability values were calculated based on the untreated cell control (Ctrl). All data shown are means ± SEM, n = 3 per group.

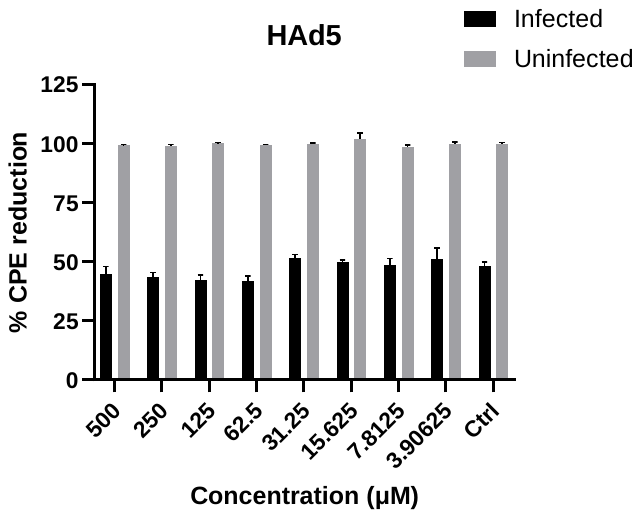

**Supplementary Fig. S2.** **Effects of teicoplanin (TP) on CPE induced by human adenovirus 5 (HAd5).** A549 cells were infected with 4,000 TCID_50_/well HAd5 and simultaneously treated with the indicated concentrations of TP. Percent cytopathic effect reduction (% CPE reduction) was calculated relative to the viability of the untreated infected and uninfected control (Ctrl). Data show means ± SEM; n = 3 per TP concentration. The graph is representative of one of at least two independent assays.
